# Chemogenetic activation of nigrostriatal dopamine neurons in freely moving common marmosets

**DOI:** 10.1101/2021.01.29.428749

**Authors:** Koki Mimura, Yuji Nagai, Ken-ichi Inoue, Jumpei Matsumoto, Yukiko Hori, Chika Sato, Kei Kimura, Takashi Okauchi, Toshiyuki Hirabayashi, Hisao Nishijo, Noriaki Yahata, Masahiko Takada, Tetsuya Suhara, Makoto Higuchi, Takafumi Minamimoto

## Abstract

To interrogate particular neuronal pathways in non-human primates under natural and stress-free conditions, we applied designer receptors exclusively activated by designer drugs (DREADDs) technology to common marmosets. We injected adeno-associated virus vectors expressing the excitatory DREADD hM3Dq into the unilateral substantia nigra in three marmosets. Using multi-tracer positron emission tomography imaging, we detected DREADD expression in vivo, which was confirmed in nigrostriatal dopamine neurons by immunohistochemistry, and assessed activation of the substantia nigra and dopamine release following agonist administration. The marmosets rotated in a contralateral direction relative to the activated side 30–90 min after consuming food containing the highly potent DREADD agonist deschloroclozapine (DCZ), but not on the following days without DCZ. These results indicate that non-invasive and reversible DREADD manipulation will extend the utility of marmoset as a primate model for linking neuronal activity and natural behavior in various contexts.

## Introduction

There has been growing interest in the common marmoset (Callithrix jacchus), a small non-human primate, as a neurobiological model of the human brain in health and disease (1),(2),(3). Owing to their rich natural behavioral repertoires with learning capabilities, marmosets have been used in a number of studies to examine the neural basis of cognitive, emotional, and social functions (4),(5). For example, a variety of functions of the nigrostriatal dopamine (DA) system have been studied in freely behaving marmosets, including cognitive flexibility (6), discrimination learning (7), and motor activity (8), through its irreversible manipulation of the DA system such as DA depletion and receptor knockdown. However, given that the individual variability is less controllable in behavioral examination, especially under freely moving conditions, reversible manipulation is more advantageous because it permits within-subject comparisons. Reversible manipulation is possible via local drug infusion (9); however, this requires that subjects are captured and held upon every manipulation, thereby causing potential adverse reactions (e.g., stress) that could confound the main effects. As such, the conventional manipulation methodologies have some limitations for the reversible and stress-free interrogation of particular neural systems, preventing neuroscientists from taking advantage of the rich natural behavior of marmosets and extending their utility as a primate model.

The chemogenetic technology designer receptors exclusively activated by designer drugs (DREADDs) affords a minimally invasive means of reversibly and remotely controlling the activity of a neuronal population expressing designer receptors through systemic delivery of their agonist (10),(11). DREADDs have widely been used to modify neuronal activity and behavior in rodents. They have also proven valuable in non-human primates, allowing reversible manipulation of activity across a specific neuronal population in large and discontinuous brain regions that are beyond the reach of pharmacological, electrical, or optogenetic manipulation strategies (12),(13),(14),(15). Because there is no requirement for chronic surgical implantation of any invasive devices, DREADDs permit flexible experimental designs to control neuronal activity in freely moving marmosets with minimal invasiveness. To date, no study has reported the application of DREADD-based neuronal control in behaving marmosets. To apply DREADDs technology to marmosets, strategies used for macaque monkeys, such as viral vector delivery and non-invasive monitoring of DREADD expression and function, should be useful and applicable. In addition, the use of the highly potent novel DREADD agonist deschloroclozapine (DCZ) should provide a wider effective window for the selective activation of DREADDs with minimal off-target effects, as demonstrated in mice and macaques16. However, conventional routes of agonist administration, such as intramuscular and intraperitoneal (IP) injections, can induce pain and stress, leading to unwanted effects on natural behavior. Because DCZ displays good brain penetrability and metabolic stability in macaques, oral delivery (per os [PO]) is a feasible alternative route for non-invasive DCZ administration for marmosets.

We therefore performed a proof-of-concept study employing non-invasive chemogenetic manipulation of a specific neural system in freely moving marmosets. We aimed to increase the excitability of nigrostriatal DA neurons, which are known to induce rotation behavior in rodents (16),(17),(18). Marmosets received injections of adeno-associated virus (AAV) vectors expressing the excitatory DREADD hM3Dq into the unilateral substantia nigra (SN). Subsequent multimodal positron emission tomography (PET) imaging was used to assess DREADD expression and function following agonist administration. We revealed that oral DCZ administration reversibly induced contralateral rotation behavior. Our results demonstrated that non-invasive chemogenetic manipulation can enhance the utility of marmoset as a primate model for linking a particular neural pathway and non-restricted behavior in a variety of contexts.

## Results

### Imaging-guided transduction and monitoring hM3Dq expression in the unilateral SN

Three marmosets received an injection of AAV vectors co-expressing hM3Dq and the green fluorescent protein (GFP) into one side of the SN under intraoperative X-ray computed tomography (CT) guidance (Table 1, Fig. 1a-b). Another control vector expressing a fluorescent protein marker (monomeric Kusabira Orange, mKO) was also injected into the opposite side. More than 6 weeks after the injection, we performed PET using the DREADD-selective radioligand [^11^C]DCZ to visualize hM3Dq expression in vivo, as demonstrated previously in macaque monkeys (15). hM3Dq expression in the SN was visualized as a region exhibiting high radioligand binding (Fig. 1c). The binding potential relative to a non-displaceable binding (BP_ND_) of [^11^C]DCZ in the target SN was significantly higher than that of the contralateral side (p = 0.038, T[1,5] = −5.017, N = 3, pairwise t-test, Fig. 1d). Postmortem immunohistochemical analysis confirmed that GFP, a marker of transgene expression, was expressed in the neurons on the target side of the SN pars compacta (SNc) in all three marmosets (e.g., Fig. 1e-f). GFP-labeled axons were also found in the ipsilateral striatum, the area receiving direct projections from the SNc (Fig. 1g-i). Thus, these results suggested that hM3Dq was introduced in the dopaminergic neurons in the SNc unilaterally in all three animals.

**Table 1.**
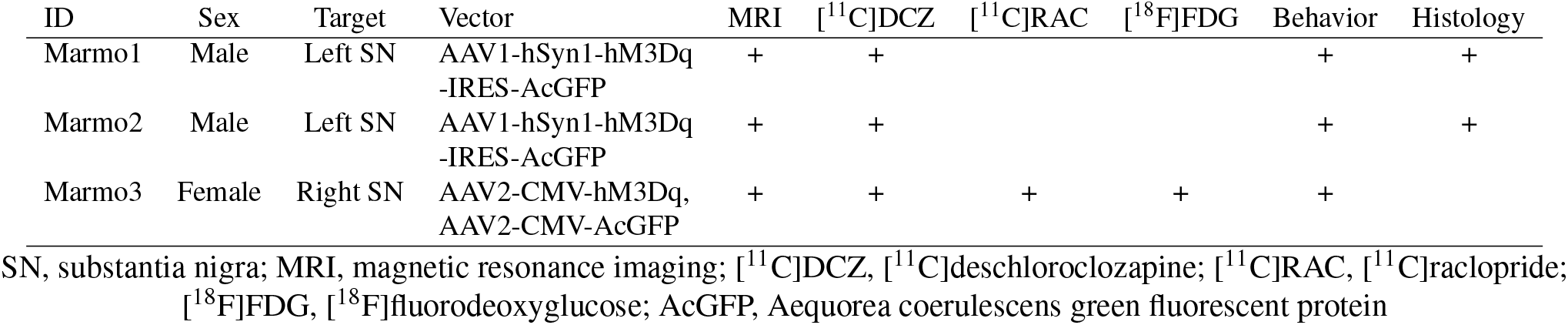
Comparison of the fitted potential energy surfaces and ab initio benchmark electronic energy calculations

**Fig. 1.**
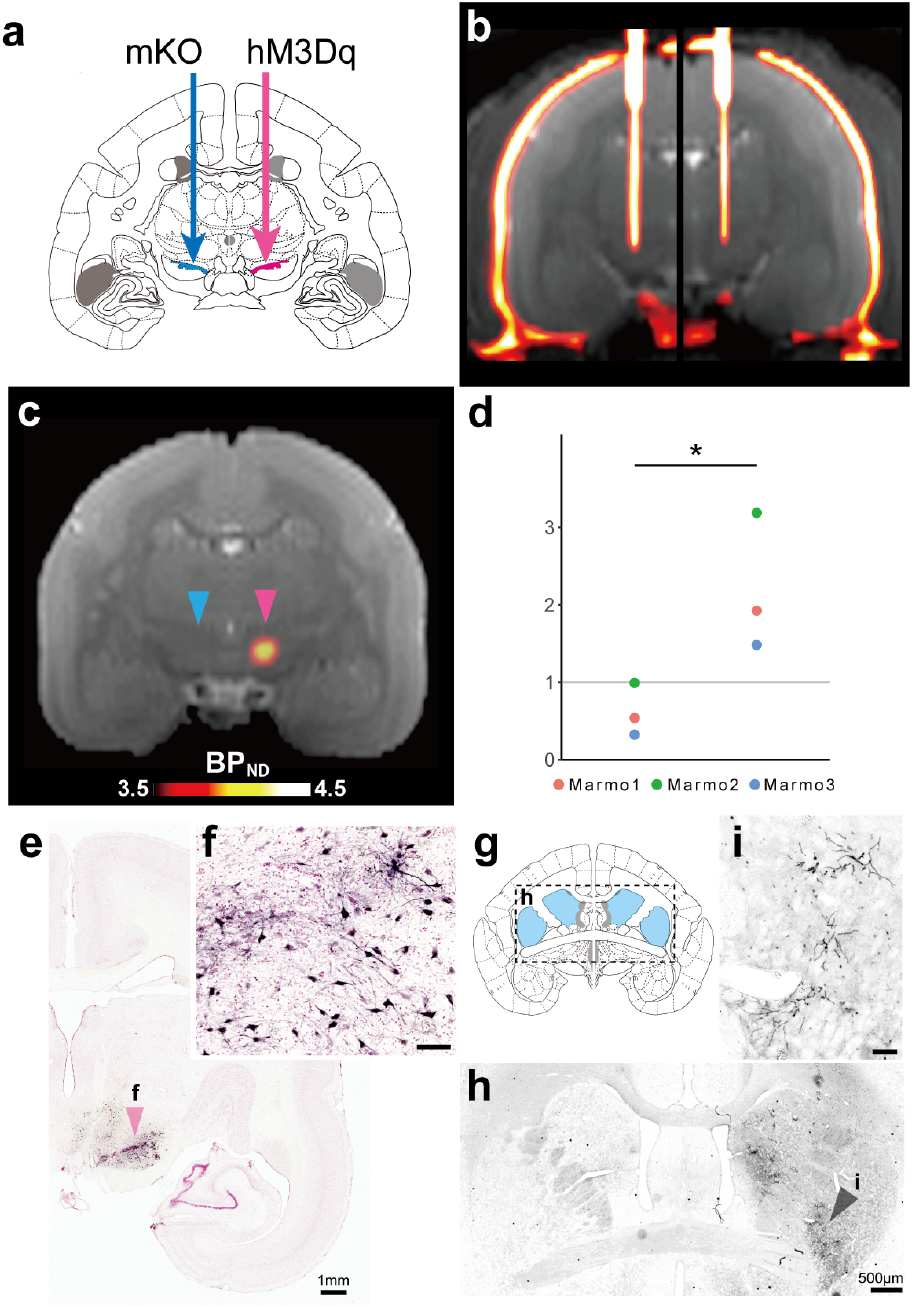
Imaging-guided injection of adeno-associated virus (AAV) vectors and evaluation of hM3Dq expression (a) Illustration of the location of viral vector injections. AAV vectors expressing hM3Dq were injected into the unilateral substantia nigra (SN, red arrow). As a control, an AAV vector (AAV2-CMV-mKO) was injected into the contralateral SN (blue arrow). (b) A coronal computed tomography image overlaid on a magnetic resonance image representing the position of the vector injection needle obtained during surgery. (c) A coronal representative parametric positron emission tomography image presenting the specific binding (BPND) of [^11^C]deschloroclozapine (DCZ; Day 43, Marmo1). Red and blue triangles denote the hM3Dq- and mKO-expressing sides, respectively. (d) Comparison of regional BPND of [^11^C]DCZ between the hM3Dq and control side of the SN for three marmosets. The asterisk indicates a significant difference (p < 0.01, pairwise t-test). (e) A photograph of the representative coronal section staining for green fluorescent protein (GFP) and neutral red including the SN (Marmo1). (f) A high-magnification image of SN neurons (red triangle in e). Scale bar = 100 μm. (g) Illustration of the location of the striatum depicted in (h) in a coronal section. (h) A coronal GFP-stained section including the striatum. (i) A high-magnification image of the striatum (black triangle in h) presenting GFP-positive axons. Scale bar = 100 μm. mKO, monomeric Kusabira Orange.

### Chemogenetic activation of the unilateral nigrostriatal DA neurons

Next, we tried to induce and monitor chemogenetic activation of the unilateral nigrostriatal neurons by hM3Dq. To monitor DREADD-induced neuronal activation in vivo, we performed PET with [^18^F]fluorodeoxyglucose (FDG) following intravenous (IV) clozapine N-oxide (CNO) administration (10 mg/kg) to detect a change in regional brain glucose metabolism, an index of brain neuronal excitability (19),(20). We observed increased uptake of [^18^F]FDG on the hM3Dq-expressing side of the SN (Fig. 2a), likely reflecting the increased metabolic activity of hM3Dq-expressing neurons. No comparable lateralized uptake was found in the other areas.To examine chemogenetically-induced DA release at postsynaptic sites, we performed PET using [^11^C]raclopride, a D2 receptor radioligand sensitive to endogenous dopamine (21),(22). We found a decrease in tracer binding specifically in the ipsilateral striatum following CNO administration (10 mg/kg IV, Fig. 2b), but not following treatment with the vehicle control, suggesting increased DA release at postsynaptic sites. Collectively, these results suggest that activation of the unilateral nigrostriatal DA neurons can be induced by an excitatory DREADD.

**Fig. 2.**
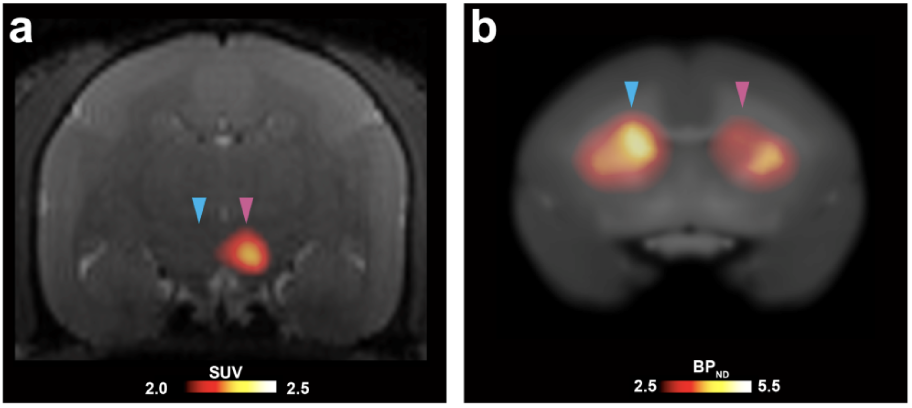
In vivo evaluation of chemogenetic activation of unilateral nigrostriatal dopaminergic (DA) neuron function (a) A coronal section of parametric image displaying the standardized uptake value (SUV) of [^18^F]fluorodeoxyglucose overlaying a magnetic resonance image following clozapine N-oxide (CNO) administration (10 mg/kg intravenous [IV]; Day 43). Red and blue triangles indicate the hM3Dq-expressing substantia nigra and its contralateral control side, respectively. (b) A coronal section of parametric image representing specific binding (BPND) of [^11^C]raclopride overlaying the magnetic resonance image following CNO administration (10 mg/kg; Day 97). Red and blue triangles indicate the ipsilateral and contralateral striatum relative to the hM3Dq-expressing side, respectively. Data were obtained from Marmo3. BP_ND_, binding potential relative to a non-displaceable radioligand.

### Minimally invasive and reversible control of unilateral DA neurons induced rotation behavior

Having successfully applied chemogenetic excitation of the unilateral nigrostriatal DA neurons, we next examined its effects on the behavior of conscious marmosets. We used the highly potent and selective DREADD agonist DCZ (15). Taking advantage of the good brain permeability of DCZ, we adopted two routes of systemic administration for behavioral experiments: IP and PO. We expected that IP delivery would provide a relatively rapid effect because of its better absorption, whereas PO deliver offers the benefit of stress and pain-free operation. Prior to behavioral experiments, we performed pharmacokinetics analysis. Following IP administration (100 μg/kg), serum DCZ concentrations rapidly increased and peaked after 26.2 ± 7.5 min, whereas PO administration (100 μg/kg) resulted in a slower increase in serum DCZ levels, which peaked after 97.5 ± 45 min (N = 4, p = 0.017, BM Statistic = 4.1, df = 3.65, Brunner-Munzel test for median values; Fig. 3). In both cases, the peak DCZ concentration (50–100 nM) was comparable or even higher than that observed in macaque monkeys following the intramuscular injection of DCZ at the same dose (15). Considering an effective dose for hM3Dq for macaques (>1 μg/kg) (15), we selected 3-10 μg/kg as the DCZ doses for subsequent behavioral experiments.

**Fig. 3.**
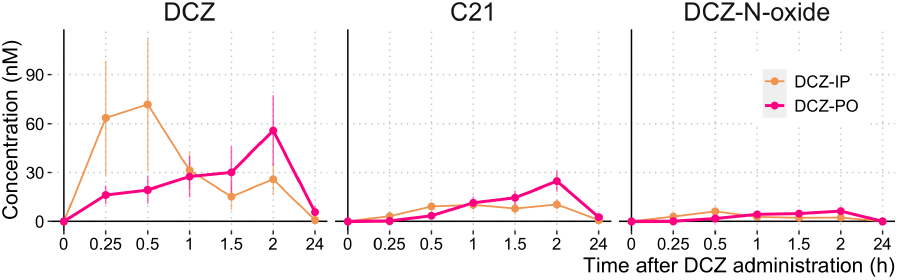
Time-concentration profiles of deschloroclozapine (DCZ) and its metabolites in plasma (DCZ 100 μg/kg per os [PO] or intraperitoneal [IP], mean ± SEM, N = 4).

We then tested the non-invasive chemogenetic control of the unilateral dopaminergic system in freely moving marmosets (Marmo1 and Marmo2; Table 1). Each marmoset was placed in a Plexiglas cylinder chamber and given a piece of food containing DCZ solution (10 μg/kg) or vehicle. All marmosets voluntarily ate the food immediately upon receipt (Movie S1). Approximately 30 min after eating the DCZ-containing food, the marmosets began to exhibit head turning, and they eventually rotated in a contralateral direction to the hM3Dq-positive side (Movie S2).

To quantify the behavioral alterations, we developed a three-dimensional (3D) motion tracking system (MTS) consisting of four depth cameras (Fig. 4a-d). Briefly, the MTS enables recording of marmoset free motion and its reconstruction as high-resolution 3D point clouds (< 3-mm isovoxel) utilizing frame-by-frame parametric physical stimulation, measuring the trajectories of body parts (*Head*, *Body*, *Neck*, and *Hip*) without body markers (Fig. 4; see Methods). MTS visualized DCZ-induced behavioral changes of the marmosets. Figure 5a presents a typical example of a top view of *Head* trajectory following vehicle and DCZ treatment (PO) and 24 h after DCZ treatment in a marmoset (Marmo2). After vehicle treatment, the marmoset exhibited “stop-and-go” behavior, as indicated by sparse ‘dotted chain’ patterns of *Head* trajectory (Fig. 5a, left). Following DCZ administration, the marmoset displayed rotation behavior, as represented by a ‘tangled ball of yarn’-like *Head* trajectory (Fig. 5a, middle). Twenty-four hours after administration, the marmoset displayed a similar movement pattern as observed in the vehicle treatment sessions (Fig. 5a right). We also found that following DCZ administration, the marmosets continuously deviated their heads contralaterally to the hM3Dq-positive side relative to the body axis. To quantify this behavioral change, we analyzed the shifts of the *Head* and *Hip* positions in every 2 s (Fig. 5b). The density plots revealed the contralateral deviation of the *Head* position and *Hip* deviation to the opposite side, which specifically occurred 30–60 min after DCZ administration (Fig. 5b, right).

**Fig. 4.**
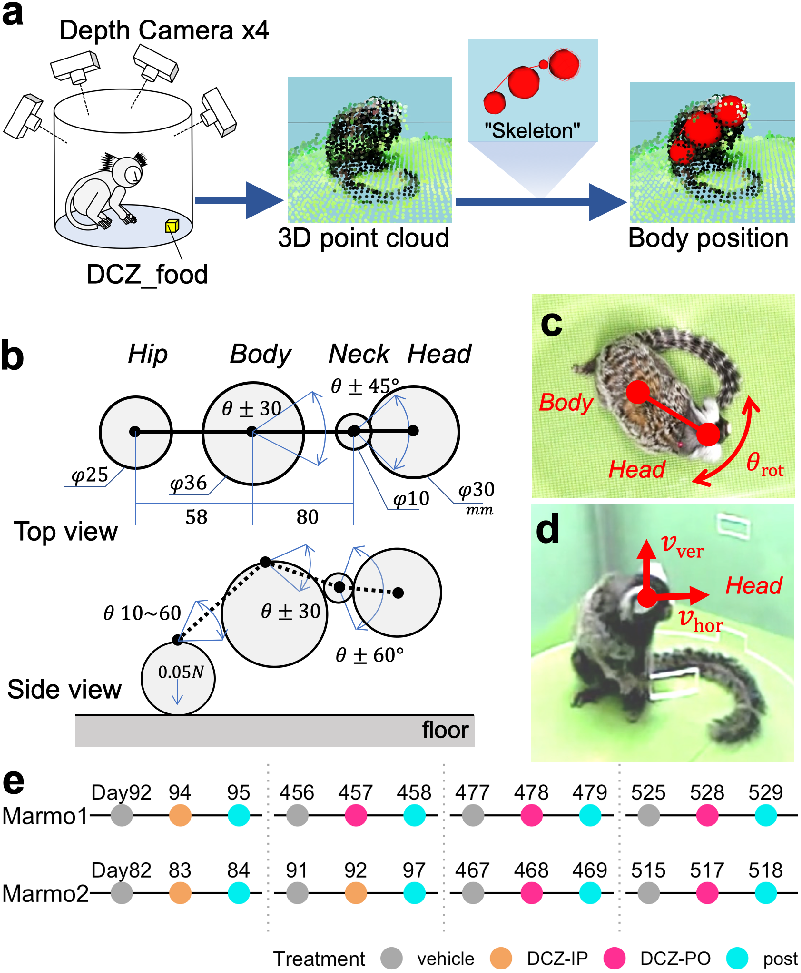
(a) Illustration of the analytical data flow of the MTS. The positions of animal body parts, namely Head, Neck, Body, and Hip, were estimated through a statistical skeleton model that simulated the full three-dimensional point cloud of the marmoset body surface extracted from four depth cameras (b) Schematics of a marmoset body skeleton model. Arrows represent the part sizes, attraction force between Hip and floor, and the range of movement of joints. (c) Body rotation speed (*θ*_rot_) was the angular speed of the horizontal Body-Head axis. (d) Head movement was divided into horizontal (V_hor_) and vertical vectors (V_ver_). (e) Experimental schedule. The marmosets were examined under conditions (vehicle treatment, deschloroclozapine [DCZ] treatment, and 24 h after DCZ treatment) using a repeated-measures design with a total of four repetitions of either per os [PO] or intraperitoneal [IP] administration. The numbers represent the days after vector injection.

**Fig. 5.**
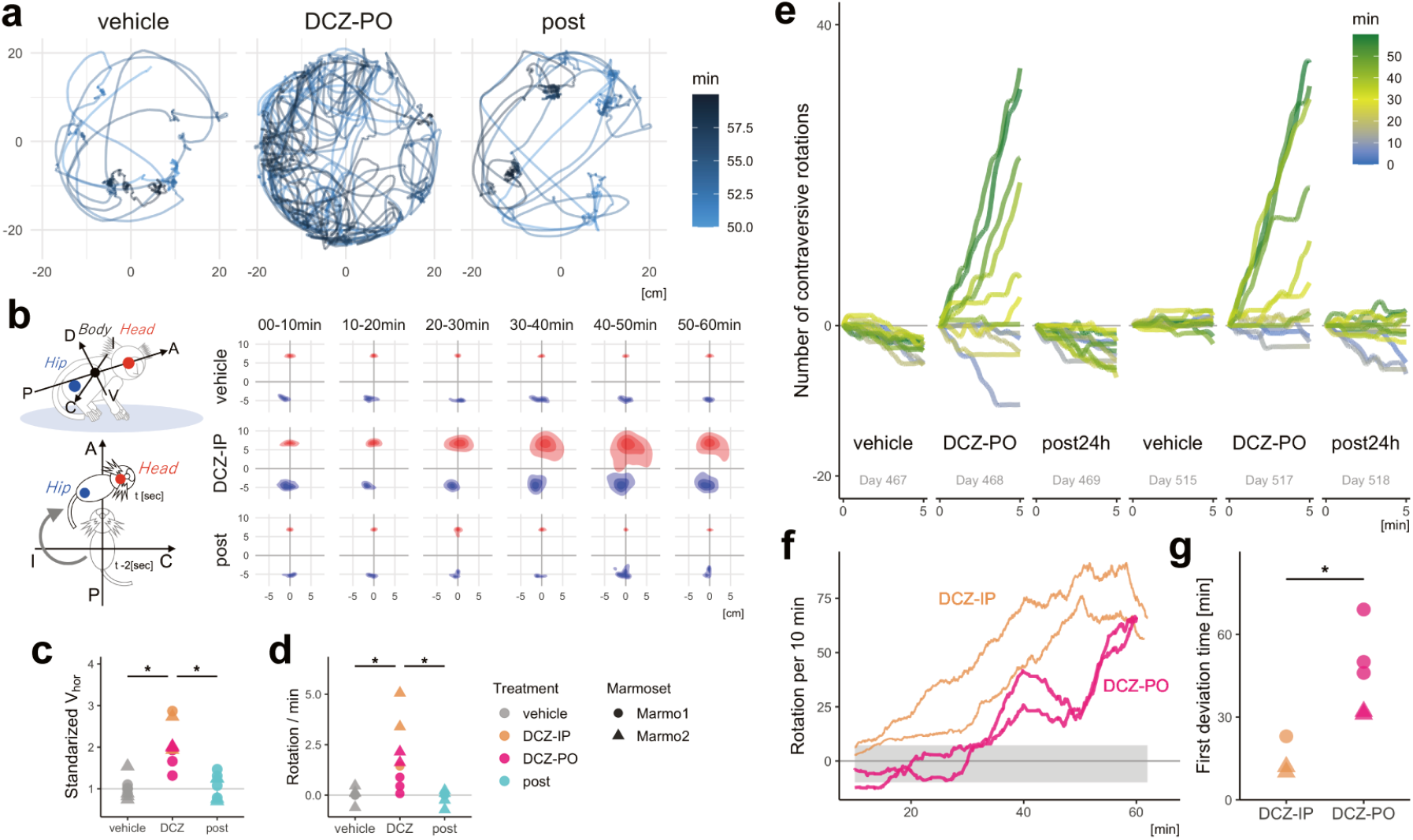
(a) Example of the top view of the Head trajectory of Marmo2 50–60 min after vehicle and deschloroclozapine (DCZ; 3 μg/kg, per os [PO]) administration and 24 h after DCZ administration. (b) Postural changes characterized by the positions of the Head and Body relative to the body axis. Left top: The body axis, i.e. anterior-posterior (A–P) axis, is defined as that through the center of the Head and Body, whereas the ipsilateral-contralateral (I–C) and dorsal-ventral (D–V) axes are defined as those horizontally and vertically orthogonal to the A–P axis. Left bottom: Relative Head and Hip positions were determined every 2 s in the A–P I–C plane. Right: Density plots of the Head (red) and Hip (blue) positions over a 10-min period following vehicle and DCZ (IP) administration and 24 h after DCZ administration. The darkness of the color indicates the quartile distribution density (75, 50, 25 percentiles) of the relative position of each body part. (c) Comparison of the horizontal Head speed (Vhor) scaled per individual with the vehicle conditions equal to 1 in animals treated with vehicle and DCZ (blue: 3 μg/kg IP; red: 10 μg/kg PO), and 24 h after DCZ administration (post) for Marmo1 and Marmo2. (d) Comparison of the frequency of contralateral rotation among three conditions. (e) Representative trends of the cumulative sum of the contraversive rotation every 5 min across conditions in Marmo2. Color scale representing the time after the treatment/test start. (f) Total number of contraversive rotations for every sliding 10-min window for IP (blue) and PO DCZ treatment (red) in Marmo2. Gray shadow represents the 95% confidence intervals of the cumulative sum of *θ*_rot_ under the vehicle condition. (g) Comparison of the latency of significant behavioral changes between IP and PO DCZ treatment, which was defined as the first deviation time from the 95% confidence interval of those under the vehicle condition. Asterisks indicate a significant difference (p < 0.01, two-way ANOVA).

Both marmosets repeatedly and selectively exhibited contralateral rotation following PO and IP administration. Both IP (3 μg/kg) and PO (10 μg/kg) DCZ administration significantly increased horizontal *Head* velocity (p = 9.8 × 10^−4^, F[2,21] = 13.64, two-way ANOVA, Fig. 5c) but not vertical velocity (p = 0.98, F[2,21] = 0.023, two-way ANOVA). We quantified the degree of marmoset body rotation by measuring the sum of rotation of the horizontal body axis (*θ*_rot_, Fig. 5d). We found that the frequency of contralateral rotation was significantly higher following DCZ administration than after vehicle administration and at 24 h after DCZ administration (DCZ vs. vehicle, p = 0.0018, diff. = 1.87; DCZ vs. post, p = 0.0012, diff. = 1.94; vehicle vs. post, p = 0.99, diff. = 0.07, Tukey’s HSD test, F[2,21] = 11.3, Fig. 5d). We also examined the temporal dynamics of the behavioral change by calculating the cumulative sum of rotation in 5-min windows. As presented in Fig. 5e, Marmo2 exhibited contralateral *Head* orientation bias 30–60 min after oral DCZ administration (yellow-green lines in Fig. 5e). This bias disappeared after 24 h but reappeared after the second DCZ dose.

We measured the latency of significant behavioral changes, which was defined as the period in which deviation of the cumulative sum of *θ*_rot_ from the 95% confidence intervals of that in vehicle-treated sessions (Fig. 5f). The latency of behavioral change was shorter in the IP sessions (15 ± 7 min) than in the PO sessions (45.6 ± 15.5 min; p = 0.004, F[1,6] = 24.3, two-way ANOVA, Fig. 5g), likely reflecting rapid absorption of the IP-delivered drug and high DCZ availability in blood (Fig. 3). Contralateral rotation was continuously observed until the end of the testing session (approximately 90 min). In contrast to the hM3Dq-expressing marmosets, non-DREADD-expressing marmosets did not display altered *Head* trajectories or *Head* deviations following DCZ administration (N = 2; Fig. S1), suggesting that the behavioral changes in Marmo1 and Marmo2 were attributable to hM3Dq activation.

## Discussion

Using DREADDs, we demonstrated that temporal activation of the unilateral nigrostriatal DA system induced contralateral rotation behavior in the common marmosets. To the best of our knowledge, this is the first study to demonstrate chemogenetic control of neuronal activity and behavioral action in common marmosets. A combination of our experimental methods employed in this study, including (1) imaging-guided viral vector delivery, (2) in vivo monitoring of DREADD expression and function, (3) painless and stress-free delivery of DREADD actuator, and (4) markerless motion tracking analysis, was critical to our successful chemogenetic manipulation and behavioral analysis in marmosets under freely moving conditions.

The specific delivery of transgenes is a critical step for successful chemogenetic manipulation, especially for deep brain structures. The use of non-invasive imaging methods, such as magnetic resonance imaging (MRI) and CT, has been proposed for the assessment of the injection site, even during surgery (23),(24). The present study used CT to visualize the injection cannula (33 gage/210 μm diameter) with 80-μm spatial resolution. Intraoperative CT was successfully used to adjust the position of the injection cannula to the target at the co-registered MR image, helping to localize hM3Dq expression at the SN. To manipulate the nigrostriatal neurons, we used two AAV vector constructs, namely AAV2-CMV-hM3Dq and AAV1-hSyn1-hM3Dq-IRES-AcGFP (Table 1), both of which confer neuron-specific expression (25),(26). Even without a DA-specific expression system, histological confirmation indicated that hM3Dq was successfully expressed in the DA neurons in the SNc. Because of the lack of specificity and a relatively large-volume injection, GFP-positive neurons were also found outside the SN. In future studies, the use of a DA neuron-specific promoter, such as a tyrosine hydroxylase promoter (27), should be taken into consideration, together with a smaller injection volume to increase the specificity of the transgene. Despite these shortcomings, our model allowed us to conduct the chemogenetic manipulation of marmoset behavior.

Prior to testing the behavioral effect of DREADD activation, it is useful to clarify designer receptor expression and function. In previous studies, PET using radiolabeled DREADD agonists was used to visualize the site and level of expression of hM4Di/hM3Dq in living rodents and macaques (14),(15),(28),(29),(30). In the present study, we performed PET using [^11^C]DCZ, a selective radioligand validated in hM3Dq-expressing mice and macaques(15). We visualized hM3Dq expression as increased tracer binding specifically at the target location in all three marmosets, which was consistent with the in vitro immunohistochemical data. We also used PET to monitor the functional effects of DREADDs. First, glucose metabolism was measured using [^18^F]FDG as an in vivo representation of neuronal activity (31). Increased [^18^F]FDG uptake was observed in hM3Dq-expressing regions following agonist administration, indicating chemogenetic activation of the SN. Second, endogenous DA release was assessed using [^11^C]raclopride, which has repeatedly been used in humans and animals, including non-human primates (21),(32). Agonist administration decreased the tracer binding on the hM3Dq-positive side of the striatum, suggesting an increase in DA release. Our data indicate that the combination of chemogenetics with non-invasive imaging methods should be valuable for future translational studies towards the development and validation of therapeutic methodologies, for example, cell replacement therapy of Parkinson’s disease (33).

When examining chemogenetic effects on behavior, it is critical to minimize the adverse side effects of agonist administration itself. Conventional administration routes such as IP may cause pain and stress in the subjects. Previous DREADD studies in rodents used non-invasive CNO delivery through drinking water or food (34),(35),(36),(37). Although the timing and dose are less precise and controllable, PO delivery of CNO has proven useful for chronic chemogenetic manipulation. In the current study, we used DCZ, a highly potent DREADD agonist with better brain permeability than CNO and C21(15). Because the effective dose of DCZ is approximately 1% of that of CNO, the solution volume (approximately 0.1 mL) was sufficiently small to permit its addition to food. Because the animals always ate the DCZ- or vehicle-containing food immediately, PO administration is an optimal non-invasive agonist delivery method for marmosets with better control of the dose and timing. Indeed, our results indicated that the PO delivery of DCZ induced behavioral changes in marmosets via hM3Dq, in line with the results following IP injections. Further, we believe that our PO protocols are also beneficial and applicable to long-lasting or chronic DREADD manipulation in marmosets.

Unlike CNO, the metabolites of which cause off-target effects, DCZ does not produce brain-permeable metabolites in mice and macaques (15). Our pharmacokinetic analysis in marmosets indicated that C21, a metabolite of DCZ, accumulated in plasma at a certain amount. Because C21 displays weak brain permeability, the effects of converted C21 on DREADDs should be negligible. Although no significant effect of DCZ (10 μg/kg PO) and its metabolites on movement speed and direction was found in non-DREADD subjects (Fig. S1), potential off-target effects should be considered in future studies in marmosets using an inhibitory DREADD, which may require higher DCZ doses.

It has repeatedly been demonstrated that contralateral rotation behavior is induced by activating the unilateral nigrostriatal system via electrical stimulation or the intrastriatal injection of DA releasers in rats (16),(17),(18). In accordance, the current study demonstrated that chemogenetic activation of unilateral nigrostriatal DA neurons induced rotation behavior contralaterally to the activated side in marmosets. As mentioned previously, chemogenetic activation was not necessarily induced in DA neurons exclusively, because hM3Dq expression was detected in some neurons outside the SNc, including SN pars reticulata (SNr) neurons. Hence, chemogenetic activation of these off-target neurons might have contributed to the observed behavioral changes. However, excitation of SNr neurons would result in ipsiversive rotation behavior (38),(39), therefore, its contribution to our model may be minor. Taken together, the most plausible explanation of the observed rotation behavior is the result of chemogenetic activation of DA neurons in SNc.

Although the behavioral alterations in our study were apparent and easily detected via visual inspection (Movie S2), objective quantification is critical for reproducibility and deep understanding of the behavioral consequences of chemogenetic manipulation. In particular, the posture of animals — a series of movements of the major body parts — reflects both motor function and their internal state such as emotion and intention (40),(41),(42),(43),(44),(45). Conventional systems available for marmosets focus on simple macroscale parameters (e.g., circadian activity rhythms (46)) or a few micro behavioral parameters (e.g., eye and face orientation (47)). We developed a markerless MTS that tracks and digitizes the 3D trajectory of the major body parts of marmosets, thereby reconstructing and analyzing the sequence of posture. Our results indicated the MTS provides a powerful means to visualize and quantify the effects of chemogenetic activation of the DA neurons in a data-driven manner.

One shortcoming of our system was the relatively high computation costs of the measurement and calculation processes, which were not fully automated but were divided into several sub-processes because of a large amount of data (approximately 20 GB per 30 min). Thus, the real-time detection of specific behavioral events remains a challenge for future work. We are currently extending this system to examine two marmosets simultaneously, making it feasible for future chemogenetic studies of social interactions.

In summary, using hM3Dq and oral DCZ administration, we manipulated the nigrostriatal DA neurons in freely behaving marmosets. We also demonstrated the utility of non-invasive imaging methods for validating DREADD expression and assessing the functional effects of DREADDs. Our results confirmed that the activity of a specific neuronal population can be controlled for hours in marmosets. A novel markerless MTS allowed analyses of multiparametric body part motion and quantification of the behavioral alterations. Collectively, these chemogenetic methods will broaden the utility of the marmoset as a primate model for clarifying the neurobiological mechanisms of emotion, cognition, and social functions and developing novel therapeutic approaches to psychiatric and neurological diseases.

## Materials and Methods

### Subjects

In total, seven laboratory-bred common marmosets were used. Three marmosets (Marmo1–3; 2.5–4.0 years old; weight, 300–400 g; Table 1) were used for the chemogenetic manipulation study, and the remaining four animals (male; 1.5–5 years old, 300-330 g) were used for pharmacokinetics analyses (N = 4) and behavioral studies as non-DREADD controls (N = 2). Each single cage was exposed to a 12-h/12-h light-dark cycle. The room temperature and humidity were maintained at 27–30°C and 40-50%, respectively. All experimental procedures were performed in accordance with the Guide for the Care and Use of Laboratory Animals (National Research Council of the US National Academy of Science) and were approved by the Animal Ethics Committee of the National Institutes for Quantum and of Radiological Sciences and Technology (#11-1038).

### Viral vector production

AAV vectors (AAV1-hSyn1-hM3Dq-IRES-AcGFP, AAV2-CMV-hM3Dq, AAV2-CMV-AcGFP, and AAV2-CMV-mKO) were produced using a helper-free triple transfection procedure and purified via affinity chromatography (GE Healthcare, USA).

### Surgery and vector injection

Surgeries were performed under aseptic conditions. We monitored body temperature, heart rate, and SpO2 throughout all surgical procedures. The marmosets were immobilized with ketamine and xylazine (10–15 and 3–5 mg/kg, intramuscularly) and then maintained under anesthesia using isoflurane (1–3%) during viral vector injection surgery. Prior to surgery, MRI and CT were performed to create overlay images to estimate the stereotaxic coordinates of target brain structures. According to the estimate, we opened burr holes (approximately 2 mm in diameter) on the skull for the injection needle. Viruses were pressure-injected using a 10-μL Nanofil Syringe (NANOFIL, WPI, USA) with a 33-gage beveled Nanofil needle (NF33BV-2, WPI). The syringe was mounted into a motorized microinjector (UMP3T-2, WPI) that was held by a manipulator (model 1460, David Kopf) on the stereotaxic frame. We obtained CT scans and fused them with prescanned MR images during surgery to verify the location of the injection needle before and after insertion (Fig. 1b). The injection needle was inserted into the brain, slowly moved down 1 mm beyond the target, and then kept stationary for 5 min, after which it was pulled up to the target location. The injection speed was set at 0.1 μL min^−1^. After each injection, the needle remained in situ for 15 min to minimize backflow along the needle. Marmo1 and Marmo2 received an injection of an AAV vector carrying the hM3Dq and Aequorea coerulescens GFP constructs (AAV1-hSyn1-hM3Dq-IRES-AcGFP; 1.0 μL; 2.0 × 10^13^ particles mL^−1^) on one side of SN and another injection of an AAV vector carrying the mKO gene (AAV2-CMV-mKO; 1.0 μL; 1.7 × 10^13^ particles mL^−1^) into the other side. Marmo3 underwent a co-injection of AAV vectors (total, 2.0 μL) carrying the hM3Dq construct (AAV2-CMV-hM3Dq; 1.2 × 10^13^ particles mL^−1^) and the AcGFP gene (AAV2-CMV-AcGFP; 0.3 × 10^13^ particles mL^−1^) into one side of SN and an injection of an AAV vector carrying the mKO gene (AAV2-CMV-mKO; 2.0 μL; 1.7 × 10^13^ particles mL^−1^) into the other side.

### Drug administration

DCZ (HY-42110, MedChemExpress, USA) and CNO (Toronto Research, Canada) were dissolved in 2.5% dimethyl sulfoxide (DMSO, FUJIFILM Wako Pure Chemical Co., Japan). These stock solutions were diluted in saline to a final volume of 0.1 mL for subsequent experimentation. For behavioral testing, DCZ was administered via IP (1 μg/kg) or PO (3 or 10 μg/kg). For pharmacokinetic studies, DCZ solution (100 μg/kg) was administered via IP or PO. For PET, CNO solution (10 mg/kg) was injected intravenously.

### PET

PET was performed using a microPET Focus 220 scanner (Siemens Medical Solutions, USA). The marmosets were immobilized via intramuscular injections of ketamine (5-10 mg/kg) and xylazine (0.2–0.5 mg/kg) and then maintained under anesthesia using isoflurane (1-3%) during all PET procedures. A transmission scan using a spiraling 68Ge point source was performed to correct attenuation before a bolus injection of the radioligands [^11^C]DCZ and [^11^C]raclopride. Emission scans were acquired in the 3D list mode with an energy window of 350-750 keV after an IV bolus injection of [^11^C]DCZ (97.3–105.4 MBq), [^11^C]raclopride (46.8–131.4 MBq), and [^18^F]FDG (88.4 MBq). For PET scans using [^11^C]raclopride and [^18^F]FDG, each animal received IV treatment with 10 mg/kg CNO or vehicle (DMSO) in advance to permit peak chemogenetic activation via hM3Dq (10 and 30 min, respectively).

### Histology

Marmosets were sedated with 25 mg/kg ketamine hydrochloride administered intramuscularly and then euthanized using an overdose of 100 mg/kg sodium pentobarbital (Somnopentyl, BCM International, Hillsborough, USA) administered intraperitoneally. Next, they were intracardially perfused with 0.1 M potassium phosphate-buffered saline (PBS), followed by 4% paraformaldehyde (Merck, Whitehouse Station, USA) in 0.1 M PBS. The brains were removed from the skull, postfixed in the same fresh fixative overnight, and saturated with 30% sucrose in PB at 4°C. A freezing microtome was used to cut coronal sections serially at a thickness of 40 μm. Every 12th section was mounted onto gelatin-coated glass slides and Nissl-stained with 1% cresyl violet. For immunoperoxidase staining, the sections were incubated for 2 days at 4°C with rabbit anti-GFP antibody (1:2,000; Thermo Fisher Scientific, USA). The sections were then incubated with biotinylated donkey anti-rabbit IgG antibody (1:1,000; Jackson Laboratories, USA), followed by treatment with an avidin-biotin-peroxidase complex kit (ABC Elite; 1:100; Vector Laboratories, USA). To visualize the antigen, the sections were reacted for 10–20 min in 0.05 M Tris-HCl buffer (pH 7.6) containing 0.04% diaminobenzene tetrahydrochloride (FUJIFILM Wako Pure Chemical Co., Japan), 0.04% NiCl_2_, and 0.002% H_2_O_2_. The reaction time was adjusted to make the density of background immunostaining almost identical throughout the cases. These sections were counterstained with 0.5% neutral red, mounted onto gelatin-coated glass slides, dehydrated, and coverslipped.

### Behavioral testing

Behavioral experiments were conducted in a sound-attenuated room (O’hara Co., Ltd., Japan; 2.4 (h) × 1.2 (w) × 1.6 (d) m^3^), which was away from the colony room. Any vocalization sounds from the colony room could not be detected in the room. The temperature was maintained at 27-31°C and 30-40% relative humidity. The internal space of the sound-attenuated room was ventilated and illuminated with fluorescent lighting. The experiments were performed between 11:00 and 16:00. Before the experimental session, each subject was transferred individually from the colony room to the experimental room in a small transport chamber (165 (h) × 140 (w) × 255 (d) mm^3^, Natsume Seisakusho, Japan) and placed in a transparent acrylic cylinder chamber (radius 0.5 m × height 0.5 m) in the sound-attenuated room (Movie S3). The subjects were allowed to adapt to the transport procedure and experimental environment for 4 consecutive days prior to behavioral testing. For 3D data acquisition, the test chamber was placed on a foothold with a green floor (1 m height), and four depth cameras (RealSense Depth Camera D435, Intel, USA) were placed around the chamber at the 3, 6, 9, and 12 o’clock positions. The cameras were connected in parallel to a PC (Windows 10, 64-bit) using USB-C cables (U3S1A01C12-050, Newnex Tech., USA; the distance was 1–5 m).

The behavioral experiment started when a marmoset was put in the chamber, and the recording of the movements lasted up to 90 min (Fig. 4e). The subject was returned to the colony room after the end of the recording session. The experiments were performed once a day for each subject.

The marmosets were examined under three conditions (vehicle treatment, DCZ treatment, and 24 h after DCZ treatment, Fig. 4e) using a repeated-measures design with a total of four repetitions of either PO or IP administration. In the PO condition, animals were fed a piece of sponge cake (approximately 2 g) containing DCZ solution (3 μg/kg) or vehicle (DMSO) on the floor of the test chamber at the start of the session (Movie S1). In the IP condition, the marmosets were injected with either DCZ solution (10 μg/kg) or vehicle immediately before the experiment began.

### Markerless 3D MTS

A novel markerless 3D MTS was developed to track and analyze the behavior of freely moving marmosets based on our previous systems designed for rats (43),(42) and macaque monkeys (45).

Our MTS software package for depth camera calibration, 3D data acquisition, and fundamental setup for physical simulation are available online (3DTracker-FAB, 3dtracker.org), and this software allowed us to robustly estimate the 3D trajectory of marmoset body parts semiautomatically as follows. First, the entire surface of a marmoset body was digitized as a 3D point cloud (Fig. 4a) based on depth images captured from the four depth cameras using the functions of Intel Realsense SDK (github.com/IntelRealSense/librealsense) and the Point Cloud library (pointclouds.org). Then, in the first frame of the video, a marmoset skeleton model was manually located near the 3D point cloud and automatically fitted by the physics-based algorithm. The skeleton model consisted of four spheres representing each body part (*Head*, *Neck*, *Body*, and *Hip*) and connected with joints rotating with different ranges corresponding to the anatomical constraints (Fig. 4b, see Movie S2). The fitting algorithm works as if it physically houses the skeleton model into the 3D point cloud of the marmoset with the aid of the Bullet Physics Library (ver. 2.8.1, bulletphysics.org). In the simulation, the three types of forces from the 3D points to the skeleton spheres were assumed to stabilize the model within the 3D point cloud, namely attraction forces, repulsive forces, and the attraction force from the *Hip* sphere toward the floor (43),(45). Finally, the resultant positions of the body skeleton spheres were recorded and used as the initial position for the next frame. The average error between the estimated *Head* positions projected onto 2D images and their ‘ground-truth’ based on manual tracking were 0.26 ± 0.04 and −0.41 ± 0.05 cm, on the X- and Y-axes, respectively. (mean ± sem, Fig. S2).

### MRI and CT

MRI and X-ray CT were performed under general anesthesia in the same manner as PET. MR T2-weighted images were obtained on a 7-tesla, 200-mm-bore MRI (BioSpec AVANCE-III, Bruker, Biospin, Germany) with a volume resonator with an 85-mm inner diameter for transmission (Bruker, Biospin) and an 8-channel phased array coil for reception (RAPID Biomedical, Germany). The obtained data were reconstructed using ParaVision 5.1 (Bruker Biospin). We used a 3D rapid acquisition enhancement sequence according to the following parameters: repetition time = 2500 ms; echo time = 8.93 ms; field of view (FOV) = 26 × 35 × 25.6 mm^3^, and matrix size = 128 × 128 × 64. CT scans were obtained using a cone-beam CT system (Accuitomo170, J. Morita Co., Japan), which was operated under the following conditions: tube voltage = 90 kVp, tube current = 5 mA, exposure time = 17.5 s, FOV = 140 (diameter) × 100 (height) mm, voxel size = 0.25 × 0.25 × 0.5 mm^3^, and a gray intensity of 16 bits. Overlay MR and CT images were created using PMOD image analysis software ver. 3.7 (PMOD Technology Ltd., Switzerland).

### Pharmacokinetics analysis

Acute blood samples were collected from a saphenous vein using a 24-gage indwelling needle (SR-OT2419C, Terumo Co., Tokyo, Japan) from four conscious marmosets immobilized on a retention holder (CL-4532, CLEA Japan Inc., Japan). Samples were collected at 15, 30, 60, 90, and 120 min and 24 h after IP or PO DCZ treatment (100 μg/kg). Plasma was separated, and samples (50 μL) were diluted in 150 μL of methanol and stored at −80°C until analysis. Samples were dissolved in 50 μL of 50% methanol followed by 20 μL of Granisetron solution (10 ng/mL, internal standard). Quantification of DCZ and its major metabolites DCZ-N-oxide and C21 was performed using LC-MS/MS as described previously (15).

### Data analysis

PET data analyses were performed using PMOD image analysis software to estimate stereotaxic coordinates of the target brain structure. The standardized radioligand uptake value was calculated as regional radioactivity (Bq/cm3) × body weight (g) / injected radioactivity (Bq) at 30–90 min after the injection. BP_ND_ was estimated using a parametric multilinear reference tissue model (MRTMo) with the cerebellum as a reference region (15). The values were compared using statistical methods with R version 3.6.1 and the formula BP_ND_ ~ side + individuals (pairwise t-test, df = [1, 2, 2], Fig. 1d), where side is the side ipsilateral or contralateral to the hM3Dq vector-injected side of SN.

Behavior data analyses were performed using R version 3.6.1 and its package tidyverse version 1.3.0 (48). The trajectory of body parts was filtered with a loess filter using the stats::loess() function with span = 1/300 and downsampled to 10 frames/s (Fig. 5a). The spatial head movement speed was calculated frame-by-frame and divided into horizontal and vertical vectors (Figs. 4c and 5b). The body axis was defined as the vector from the Body center to the Head center, and *θ*_rot_ was calculated from its horizontal component with a positive value for the contralateral direction to the activated side (Fig. 4d). To quantify the biased rotation behavior, the number of total contraversive rotations was calculated from the cumulative sums of *θ*_rot_ as follows:

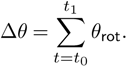

The number of contraversive rotations in 5 min (Fig. 5e) was defined as Δ*θ*_1_/2*π*, where 1 was cumulative sum in the time window [*t*_0_, *t*_1_] as follows:

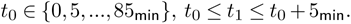

The number of contraversive rotations in 10 min (Fig. 5f) was defined as Δ*θ*_2_/2*π*, where Δ*θ*_2_ was cumulative sum in the sliding time window [*t*_0_, *t*_1_] as follows:

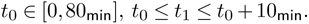

To quantify the effects of DCZ treatment, the deviation score R_in_ was calculated from 2 as the ratio of staying time under the 95% confidence interval of 2 in the vehicle condition (Fig. 5g) as follows:

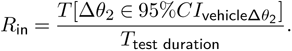

These parameters were compared between the treatment conditions across individuals as value ~ individuals + treatment for two-way ANOVA and Tukey’s honestly significant difference test.

## ACKNOWLEDGEMENTS

We thank J. Kamei, R. Yamaguchi, Y. Matsuda, Y. Sugii, T. Kokufuta, A. Maruyama,Y. Iwasawa, A. Tanizawa, S. Shibata, N. Nitta, M. Nakano and M. Fujiwara for their technical assistance. We also thank Dr. M-R. Zhang and his colleagues at Department of Radiopharmaceuticals Development, NIRS/QST for producing the radioligands. This study was supported by MEXT/JSPS KAKENHI Grant Numbers JP18H04037 (to TM), JP17H06040, JP19H04996 (to KM), and JP17H02219 (TH), and AMED under Grant Numbers JP20dm0107146 (to TM), JP20dm0307007 (to TH), JP20dm0207003 (to MT), JP20dm0307026 (to CS), JP20dm0207072(to MH), and JP20dm0307021 (to KI).

## Notes

### Competing Interest Statement

The authors have declared no competing interest.

