## Supplementary Fig. 1 for "Chemogenetic activation of nigrostriatal dopamine neurons in freely moving common marmosets"

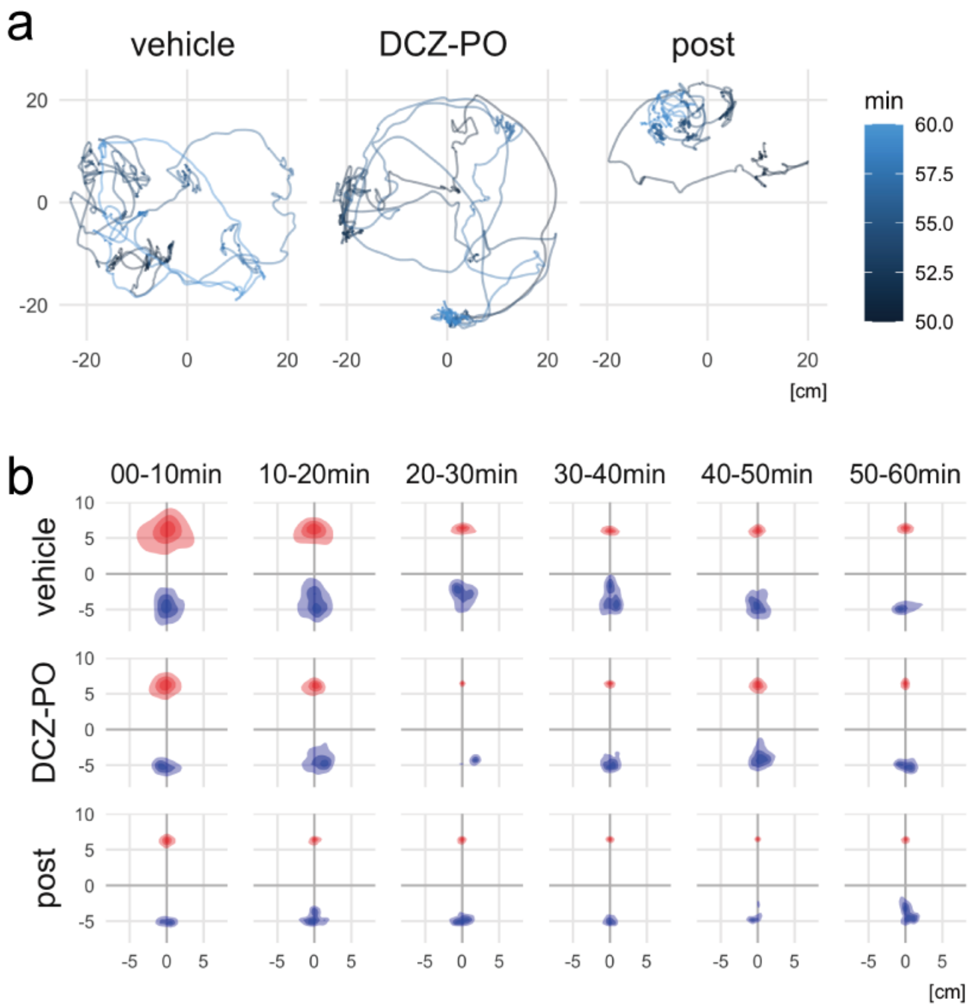


**Figure S1:** (a) Example of *Head* trajectories of a non-DREADD marmoset 50–60 min following vehicle and deschloroclozapine (DCZ) administration (10 µg/kg, *per os*) and 24 h after DCZ treatment (post). There was no significant difference in moving velocity among the treatment conditions in two non-DREADD marmosets ($p = 0.95$, $F[2, 3] = 0.057$). (b) Density plots of the *Head* (red) and *Hip* (blue) positions during a 10-min period following vehicle and DCZ administration and 24 h after DCZ administration in a non-DREADD marmoset. Unlike the case in Marmo2 shown in Fig. 4b, there was no significant deviation of *Head* or *Hip* position. DREADD, designer receptor exclusively activated by designer drugs.
