## Supplementary Fig. 2 for "Chemogenetic activation of nigrostriatal dopamine neurons in freely moving common marmosets"

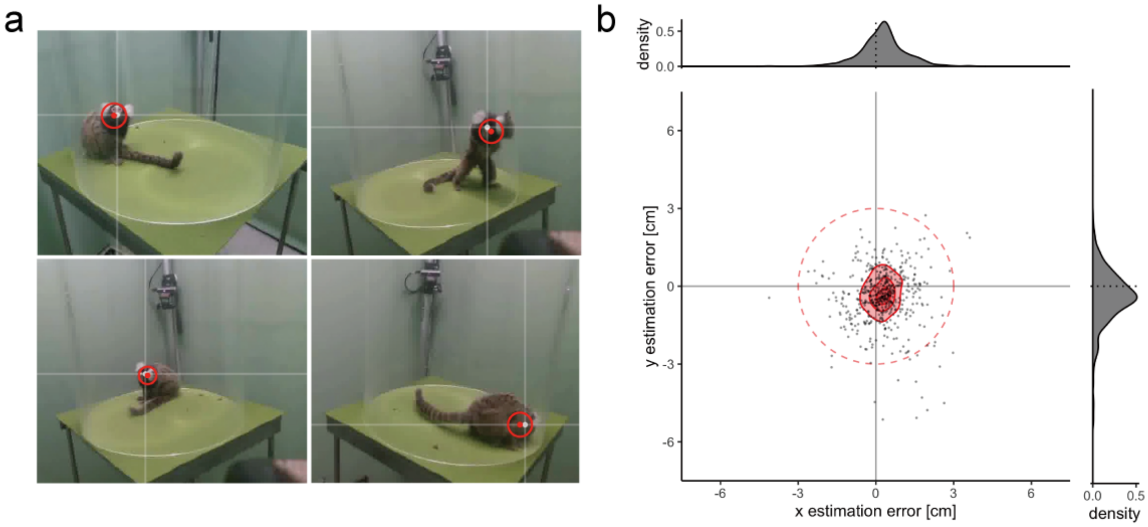


**Figure S2:** Estimated errors of the marmoset *Head* position
(a) Examples of 2D projection images of the 3D *Head* position (red circle with center point) estimated by the motion tracking system [MTS] and manually annotated head position (white point with x and y axis). (b) Scatter and density plots of error between manual annotation and MTS estimated results on 2D projection images calibrated with *Head* skeleton radius (3 cm, red circle in a) in randomly selected 500 frames. The red areas on the 2D density plot represent quantile kernel densities (75, 50, and 25 percentiles). The mean ± SEM errors were 0.26 ± 0.04 and –0.41 ± 0.05 cm, on the X- and Y-axes, respectively. 98% of estimated positions were inside of the *Head* area (distance < 3 cm).
